# Base-pair Ambiguity and the Kinetics of RNA Folding

**DOI:** 10.1101/329698

**Authors:** Guangyao Zhou, Jackson Loper, Stuart Geman

## Abstract

**Background:** A folding RNA molecule encounters multiple opportunities to form non-native yet energetically favorable pairings of nucleotide sequences. Given this forbidding free-energy landscape, mechanisms have evolved that contribute to a directed and efficient folding process, including catalytic proteins and error-detecting chaperones. Among structural RNA molecules we make a distinction between “bound” molecules, which are active as part of ribonucleoprotein (RNP) complexes, and “unbound,” with physiological functions performed without necessarily being bound in RNP complexes. We hypothesized that unbound molecules, lacking the partnering structure of a protein, would be more vulnerable than bound molecules to kinetic traps that compete with native stem structures. We defined an “ambiguity index”—a normalized function of the primary and secondary structure of an individual molecule that measures the number of kinetic traps available to nucleotide sequences that are paired in the native structure, presuming that unbound molecules would have lower indexes. The ambiguity index depends on the purported secondary structure, and was computed under both the comparative (“gold standard”) and an equilibrium-based prediction which approximates the minimum free energy (MFE) structure. Arguing that kinetically accessible metastable structures might be more biologically relevant than thermodynamic equilibrium structures, we also hypothesized that MFE-derived ambiguities would be less effective in separating bound and unbound molecules.

**Results:** We have introduced an intuitive and easily computed function of primary and secondary structures that measures the availability of complementary sequences that could disrupt the formation of native stems on a given molecule—an ambiguity index. Using comparative secondary structures, the ambiguity index is systematically smaller among unbound than bound molecules, as expected. Furthermore, the effect is lost when the presumably more accurate comparative structure is replaced instead by the MFE structure.

**Conclusions:** A statistical analysis of the relationship between the primary and secondary structures of non-coding RNA molecules suggests that stem-disrupting kinetic traps are substantially less prevalent in molecules not participating in RNP complexes. In that this distinction is apparent under the comparative but not the MFE secondary structure, the results highlight a possible deficiency in structure predictions when based upon assumptions of thermodynamic equilibrium.

## Background

Discoveries in recent decades have established a wide range of biological roles served by RNA molecules, in addition to their better-known role as carriers of the coded messages that direct ribosomes to construct specific proteins. Non-coding RNA molecules participate in gene regulation, DNA and RNA repair, splicing and self-splicing, catalysis, protein synthesis, and intracellular transportation [1, 2]. The precursors to these actions include a multitude of processes through which primary structures are transformed into stable or metastable secondary and tertiary structures. There are many gaps in our knowledge, but accumulating evidence (cf. [3–8]) suggests that the full story typically includes cotranscriptional explorations of secondary and tertiary structures, possibly accompanied by finely regulated transcription speeds, as well as a selection of proteins that may participate as stabilizers, catalysts, partners in a ribonculeoprotein complex, or chaperones to guide the process and detect errors. It is not surprising, then, that although many non-coding RNA molecules can be coxed into folding, properly, in artificial environments, the results rarely if ever match *in vivo* production in terms of speed or yield [3, 4, 9, 10].

Nevertheless, given the infamously rugged free-energy landscape of all but the smallest RNA molecules, there is good reason to expect that many of the large structural RNA molecules evolved not only towards a useful tertiary structure but also, at the same time, to help navigate the energy landscape. We reasoned that this process, a kind of co-evolution of pathway and structure, might have left a statistical signature, or “tell,” in the relationships between primary and native secondary structures. The primary structures of RNA molecules typically afford many opportunities to form short or medium-length stems,^[1]^ most of which do not participate in the native structure. This not only makes it hard for the computational biologist to accurately predict secondary structure, but might equally challenge the biological process to avoid these kinetic traps. Once formed, they require a large amount of energy (not to mention time) to be unformed.

Taking this kinetic point of view a step further, we conjectured that evolutionary pressures would tend to suppress the relative prevalence of ambiguous pairings, meaning available complementary subsequences, more for those subsequences that include paired nucleotides in the native structure than for equally long subsequences that do not. The idea being that ambiguities of stem-participating subsequences would directly compete with native stem formations and therefore be more likely to inhibit folding. Here, we do not mean to suggest that these particular adaptive mechanisms would obviate the need or advantages of other adaptations, including the reliance on proteins as both nonspecific and specific cofactors. Herschlag [3] (and many others since) argued convincingly that thermodynamic considerations applied to an unaccompanied RNA molecule could explain neither the folding process nor the stability of the folded product, explicitly anticipating multiple roles for protein cofactors. It is by now apparent that many mechanisms have evolved, and are still evolving, to support repeatable and efficient RNA folding. We are suggesting that some of these, perhaps among the earliest, might be visible upon close examination of relationships between the availability of ambiguous pairings for stem structures to those for non-stem structures. Shortly, we will introduce a formal definition of this *relative* ambiguity, which will be a molecule-by-molecule *difference* between the average ambiguity counts in and around native-structure stems and the average counts from elsewhere on the molecule. For now, we note that this measure, which we will call the *ambiguity index* and label *d*, depends on both the primary (“*p*”) and native secondary (“*s*”) structures of the molecule, which we emphasize by writing *d*(*p, s*) rather than simply *d*.^[2]^ To the extent that for any given native structure there is evolutionary pressure to minimize relative stem ambiguities, we expect to find small values of the ambiguity indexes.

But it would be a mistake to apply this line of thinking indiscriminately. The pathway to function for the many RNA molecules that operate as part of a larger, composite, complex of both RNA and protein components—the ribonucleoproteins, is considerably more complicated. The assembly of these complexes is far from fully worked out, but it stands to reason that the structures and folding of the component RNA molecules are influenced by the conformations of the accompanying proteins [8]. In such cases, the folding kinetics of the RNA molecule, as it might proceed in isolation and based only on thermodynamics and the free-energy landscape, may have little relevance to the *in vivo* assembly and arrival at a tertiary structure. Hence we will make a distinction between RNA molecules that are components of ribonucleoproteins (which we will refer to as “bound” RNA molecules) and RNA molecules which can function without being bound in a ribonucleoprotein complex (which we will refer to as “unbound” RNA molecules). The distinction is more relative than absolute. For example, many of the Group II introns both self-splice and reverse-splice, and both processes involve protein cofactors, some of which include a tight ribonculeoprotein complex with the maturase protein [7]. Nevertheless, we will treat these (as well as the Group I introns) as examples of “unbound,” since most, if not all, can function without being bound to a specific protein [10], and since there is evidence that the adaptation of preexisting proteins to function in the splicing process evolved relatively recently [11].

The advantage of the two categories, bound and unbound, is that we can avoid making difficult absolute statements about the values of ambiguity indexes, *per se*, and instead focus on comparisons across the two populations. We reasoned that molecules from the bound (ribonculeoprotein) families would be less sensitive to the kinetic traps arising from ambiguities of their stem-producing subsequences than molecules from the unbound families. We therefore expected to find smaller ambiguity indexes in the unbound families. Recall now that the ambiguity index depends on both the primary and native secondary structures of the molecule, *d* = *d*(*p, s*), which raises the question—which secondary structure *s* should be used in the calculation? Our main conclusions were drawn using the comparative secondary structure [12, 13], widely considered the gold standard.

But this dependency on *s* also afforded us the opportunity to make comparisons to a second, much-studied, approach to secondary structure prediction: equilibrium thermodynamics. The premise, namely that the structures of non-coding RNA molecules *in vivo* are in thermal equilibrium, is controversial. Nevertheless, variations on equilibrium methods constitute the prevailing *computational* approaches to predicting secondary structure.^[3]^ Typically, these approaches use estimates of the conformation-dependent contributions to the free-energy and dynamic-programming type calculations to produce either samples from the resulting equilibrium distribution or minimum free energy (MFE) secondary structures [14, 15]. Yet the biological relevance of equilibrium and minimum energy structures has been a source of misgivings at least since 1969, when Levinthal pointed out that the time required to equilibrate might be too long by many orders of magnitude [16]. In light of these observations, and considering the “frustrated” nature of the folding landscape, many have argued that when it comes to structure prediction for macromolecules, kinetic accessibility is more relevant than equilibrium thermodynamics [16–20]. In fact, a metastable state that is sufficiently long-lived and accessible might be biologically indistinguishable from an equilibrium state. Since the same issues of kinetic accessibility and the roles of kinetic traps that are behind these controversies are also behind our motivation to explore ambiguities, we also used the MFE secondary structure *s*′, as estimated using standard packages, to compute a second ambiguity index for each RNA molecule: *d*(*p, s*′). In this way, we could look for differences, if any, between conclusions based on the comparative structure and those based on the MFE structure.

The choice of RNA families to represent the two groups was limited by the availability of comparative secondary structures and the belief that the ambiguities captured by our index would be more relevant in large rather than small RNA molecules. With these considerations in mind, we chose the transfer-messenger RNAs (tmRNA), the RNAs of signal recognition particles (SRP RNA), the ribonuclease P family (RNase P), and the 16s and 23s ribosomal RNAs (16s and 23s rRNA) as representatives of “bound” (ribonucleoprotein) RNA molecules, and the Group I and Group II introns (sometimes referred to as self-splicing introns) as representatives of “unbound” molecules. See *Methods* for more details about the data set.

In summary, we will make a statistical investigation of the ambiguity index, as it varies between two groups of molecules (bound and unbound) and as it is defined according to either of two approaches to secondary structure prediction (comparative and MFE). In line with expectations, we will demonstrate that unbound molecules have systematically lower ambiguity indexes, when computed using comparative secondary structures, than bound molecules. The effect is strong: the average ambiguity in each unbound family is lower than the average ambiguity in every bound family. And the effect is still visible at the single-molecule level: a randomly chosen molecule can be accurately classified as belonging to the unbound group versus the bound group by simply thresholding on the ambiguity index (ROC area 0.81). We will also show that the utility of the ambiguity index to distinguish unbound from bound molecules disappears when the MFE structure is substituted for the comparative structure in computing the index. A related observation is that the ambiguity index of an unbound molecule can be used to classify whether the index itself was derived from the comparative versus MFE structure. To the extent that the comparative secondary structures are more accurate, these latter results might be interpreted as adding to existing concerns about the relevance of equilibrium RNA structures.

By using comparisons as opposed to absolute statistics, and various normalizations, and by favoring non-parametric (distribution-free) statistical methods, we have done our best to avoid subtle biases and hidden assumptions that would explain or at least influence the results. But more confidence would come with more data, especially more RNA families of both the ribonucleoprotein type and those that typically function without first forming tight assemblies with proteins. Given the rate of new discoveries and the rapid growth of accessible data sets, opportunities can not be far away.

The remainder of the paper is organized as follows: In the *Results* section we first develop some basic notation and definitions, and then present an exploratory and largely informal statistical analysis. This is followed by formal results comparing ambiguities in molecules drawn from the unbound families to those from the bound families, and then by a comparison of the ambiguities implied by secondary structures derived from comparative analyses to those derived through minimization of free energy. The *Results* section is followed by *Discussion and Conclusions*, in which we will recap the main results, further speculate about their interpretations, suggest refinements in the index that might highlight the effects of cotranscriptional folding, and review how our results bear on current thinking about RNA folding and structure. And finally, in *Methods*, we include detailed information about the data and its (open) source, as well as links to code that can be used to reproduce our results or for further experimentation.

## Results

### Basic Notation and the Ambiguity Index

Consider a non-coding RNA molecule with *N* nucleotides. Counting from 5′ to 3′, we denote the primary structure by

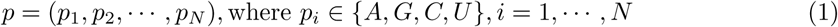

and the secondary structure by

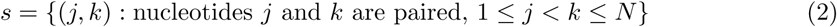

and we define the *segment* at *location i* to be

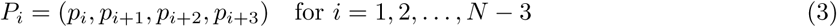

In other words, the segment at location *i* is the sequence of four consecutive nucleotides that starts at *i* and proceeds from 5′ to 3′. There is no particular reason for using segments of length four, and in fact all qualitative conclusions are identical with segment lengths three, four, or five, and quite likely, many other larger lengths.

Which segments are viable complementary pairs to *P*_*i*_, based just on location and not nucleotide content? The only constraint on location is that an RNA molecule cannot form a loop of two or fewer nucleotides. Let *A*_*i*_ be the set of all segments that are potential pairs of *P*_*i*_:

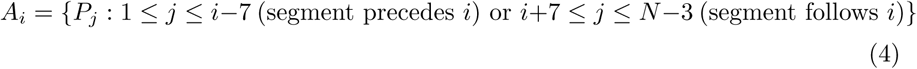

We can now define the *local ambiguity function*,

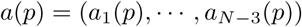

which is a vector-valued function of the primary structure *p*. The vector has one component, *a*_*i*_(*p*), for each segment *P*_*i*_, namely the number of feasible segments that are complementary to *P*_*i*_ (allowing for G·U wobble pairings in addition to Watson-Crick pairings):

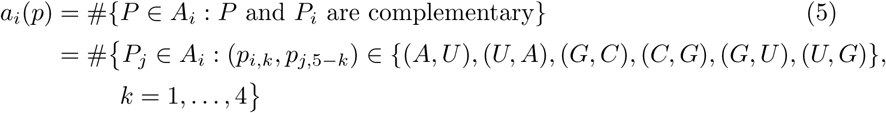

Notice that *a*_*i*_(*p*) is independent of secondary structure *s*. It is simply the total number subsequences that could form a stem structure with (*p*_*i*_, *p*_*i*+1_, *p*_*i*+2_, *p*_*i*+3_).

We want to explore the relationship between ambiguity and secondary structure. We can do this conveniently, on a molecule-by-molecule basis, by introducing another vector-valued function, this time depending only on a purported secondary structure. Specifically, the new function assigns a descriptive label to each location (i.e. each nucleotide), determined by whether the segment at the given location is fully paired, partially paired, or fully unpaired.

Formally, given a secondary structure *s*, as defined in Eq (2)2, and a location *i ∈ {*1, 2, *…, N −* 3*}*, let *f*_*i*_(*s*) be the number of nucleotides in *P*_*i*_ that are paired under *s*:

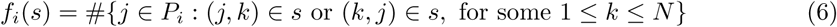

Evidently, 0 *≤ f*_*i*_(*s*) *≤* 4. The “paired nucleotides function” is then the vector-valued function of secondary structure defined as *f* (*s*) = (*f*_1_(*s*), *…, f*_*N−*3_(*s*)). Finally, we use *f* to distinguish three types of locations (and hence three types of segments): location *i* will be labeled

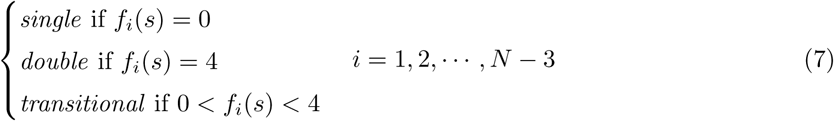

In words, given a secondary structure, location *i* is *single* if none of the four nucleotides (*p*_*i*_, *p*_*i*+1_, *p*_*i*+2_, *p*_*i*+3_) are paired, *double* if all four are paired, and *transitional* if 1, 2, or 3 are paired.

### A First Look at the Data: Shuffling Nucleotides

Our goals are to explore connections between ambiguities and basic characteristics of RNA families, as well as the changes in these relationships, if any, when using comparative as opposed to MFE secondary structures. For each molecule and each location *i*, the segment at *i* has been assigned a “local ambiguity” *a*_*i*_(*p*) that depends only on the primary structure, and a label (*single, double*, or *transitional*) that depends only on the secondary structure. Since the local ambiguity, by itself, is strongly dependent on the length of the molecule, and possibly on other intrinsic properties, we define a *relative* ambiguity index: “*d*_*T−S*_ (*p, s*)” which depends on both the primary (*p*) and purported secondary (*s*) structures:

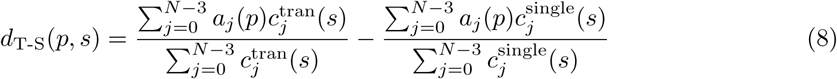

where we have used 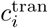 and 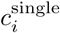 for indicating whether location *i* is *transitional* or *single* respectively. In other words, for each *i* = 1, 2, *…, N −* 3

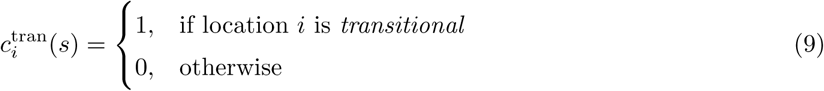

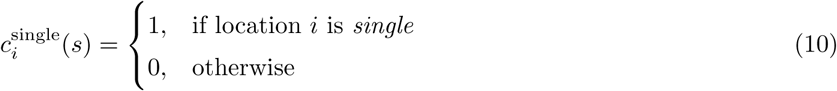

In short, the T-S ambiguity index is the difference in the averages of the local ambiguities at *transitional* sites and *single* sites.

We have also experimented with a second, closely related, index *d*_*D−S*_ (*p, s*), in which averages over *double* locations replace averages over *transitional* locations. Since the definition is somewhat complicated by the observation that local ambiguities at *double* locations are almost always greater than one (the exceptions being certain configurations with bulges), and since the results using *d*_*D−S*_ mirror those using *d*_*T−S*_ (albeit somewhat weaker), we will focus exclusively *d*_*T−S*_. Results using *d*_*D−S*_ can be accessed along with data and code, as explained in the *Methods* section. (Since there is only one index we could write *d* in place of *d*_*T−S*_, but chose to retain the subscript as a reminder of the source.)

Thinking kinetically, we might expect to find relatively small values of *d*_T-S_, at least for molecules in the unbound families, as discussed in *Background*. One way to look at this is that larger numbers of partial matches for a given sequence in or around a stem would likely interfere with the *nucleation* of the native stem structure, and nucleation appears to be a critical and perhaps even rate-limiting step. Indeed, the experimental literature [21–24] has long suggested that stem formation in RNA molecules is a two-step process. When forming a stem, there is usually a slow nucleation step, resulting in a few consecutive base pairs at a nucleation point, followed by a fast zipping step. It is important to note, though, that the application of this line of reasoning to the *d*_*T−S*_ (*p, s*) index requires that *s* be an accurate representation of the native secondary structure. For the time being we will use the time-honored comparative structures for *s*, returning later to the questions about MFE structures raised in *Background*.

How are we to gauge *d*_T-S_ and compare values across different RNA families? Consider the following experimen: for a given RNA molecule we create a “surrogate” which has the same nucleotides, and in fact the same counts of *all* four-tuple segments as the original molecule, but is otherwise ordered randomly. If ACCU appeared eight times in the original molecule, then it appears eight times in the surrogate, and the same can be said of all sequences of four successive nucleotides—the frequency of each of the 4^4^ possible segments is preserved in the surrogate. If we also preserve the locations of the *transitional, double*, and *single* labels (even though there is no actual secondary structure of the surrogate), then we can compute a new value for *d*_T-S_, say 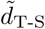, from the surrogate. If we produce many surrogate sequences then we will get a sampling of 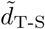 values, one for each surrogate, to which we can compare *d*_T-S_. We made several experiments of this type—one for each of the seven RNA families (Group I and Group II Introns, tmRNA, SRP RNA, RNase P, and 16s and 23s rRNA).

To make this precise, consider an RNA molecule with primary structure *p* and comparative secondary structure *s*. Construct a segment “histogram function,” ℋ (*p*), which outputs the number of times that each of the 4^4^ possible segments appears in *p*. Let 𝒫 (*p*) be the set of all permutations of the ordering of nucleotides in *p*, and let *ε* (*p*) ⊆𝒫 (*p*) be the subset of permutations that preserve the frequencies of four-tuples:

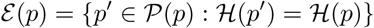

Clever algorithms exist for efficiently drawing independent samples from the uniform distribution on *ε* —see [25–27]. Let *p*^(1)^, *…, *p**^(*K*)^ be *K* such samples, and let *d*_T-S_ (*p*^(1)^, *s), …, d*_T-S_ (*p* ^(*K*)^, *s)* be the corresponding T-S ambiguity indexes. The results are not very sensitive to *K*, nor to the particular sample, provided that *K* is large enough. We used *K* =10,000. Finally, let *α*_T-S_(**p*, s*) *∈* [0, 1] be the left-tail empirical probability of choosing an ambiguity index less than or equal to *d*_T-S_(**p*, s*) from the ensemble of values {*d*_T-S_(**p*, s*), *d*_T-S_(*p*^(1)^, *s*), *…, d*_T-S_(*p*^(*K*)^, *s*):

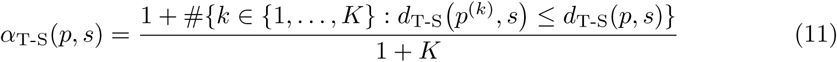

In essence, for each RNA family the *α* score is a self-calibrated ambiguity index. It is tempting to interpret *α*_T-S_(*p, s*) as a p-value from a conditional hypothesis test: Given *s* and *H*, test the null hypothesis that *d*_T-S_(*p, s*) is statistically indistinguishable from *d*_T-S_(*p*′, *s*), where *p*′ is a random sample from ℋ. If the alternative hypothesis were that *d*_T-S_(*p, s*) is too small to be consistent with the null, then the null is rejected in favor of the alternative with probability *α*_T-S_(*p, s*). The problem with this interpretation is that this null hypothesis violates the observation that given ℋ there is information in *s* about *p*, whereas *p*^(1)^, *…, p*^(*K*)^ are independent of *s* given ℋ. In other words, *d*_T-S_(*p, s*) and *d*_T-S_(*p*′, *s*) have different conditional distributions given *s* and ℋ, in direct contradiction to the null hypothesis. A larger problem is that there is no reason to believe the alternative; we are more interested in *relative* than absolute ambiguity indexes. Thinking of *α*_T-S_(*p, s*) as a calibrated intra-molecular index, we want to know how *α*_T-S_(*p, s*) varies across RNA families, and whether these variations depend on the differences between comparative and MFE structures.

Nevertheless, *α*_T-S_(*p, s*) is a useful statistic for exploratory analysis. Table 1 provides summary data about the *α* scores for each of the seven RNA families. For each molecule in each family we use the primary structure and the comparative secondary structure, and *K* =10,000 samples from *ε*, to compute individual T-S scores (Eq 11). Keeping in mind that a smaller value of *α* represents a smaller calibrated value of the corresponding ambiguity index *d*(*p, s*), there is evidently a disparity between ambiguity indexes of RNA molecules that form ribonucleoproteins and those that are already active without forming a ribonculeoprotein complex. As a group, unbound molecules have systematically lower ambiguity indexes. As already noted, this observation is consistent with, and in fact anticipated by, a kinetic point of view. Shortly, we will further support this observation with ROC curves and rigorous hypothesis tests.

**Table 1.** Comparative Secondary Structures: calibrated ambiguity indexes, by RNA family. The number of molecules, the median length (number of nucleotides), and the median *α* scores for the T-S ambiguity indexes (Eq 11) for each of the seven RNA families studied. RNA molecules from the first two families (unbound) are active without necessarily forming ribonucleoprotein complexes; the remaining five are bound in ribonucleoproteins. Molecules from the unbound families have lower ambiguity indexes.

| family | number molecules | median length | median $\alpha_{T-S}$ |
| --- | --- | --- | --- |
| Group I Introns | 116 | 451 | 0.432 |
| Group II Introns | 34 | 990 | 0.181 |
| tmRNA | 404 | 363 | 0.926 |
| SRP RNA | 346 | 274 | 0.790 |
| RNase P | 407 | 330 | 0.925 |
| 16s rRNA | 279 | 1512 | 0.938 |
| 23s rRNA | 59 | 2913 | 1.000 |

Does the MFE structure similarly separate single-entity RNA molecules from those that form ribonucleoproteins? A convenient way to explore this question is to recalculate and recalibrate the ambiguity indexes of each molecule in each of the seven families, but using the MFE in place of the comparative secondary structures. The results are summarized in Table 2. By comparison to the results shown from Table 1, the separation of unbound from bound molecules nearly disappears when viewed under the MFE secondary structures. Possibly, the comparative structures, as opposed to the MFE structures, better anticipate the need to avoid kinetic traps in the folding landscape. Here too we will soon revisit the data using ROC curves and proper hypothesis tests.

**Table 2.**
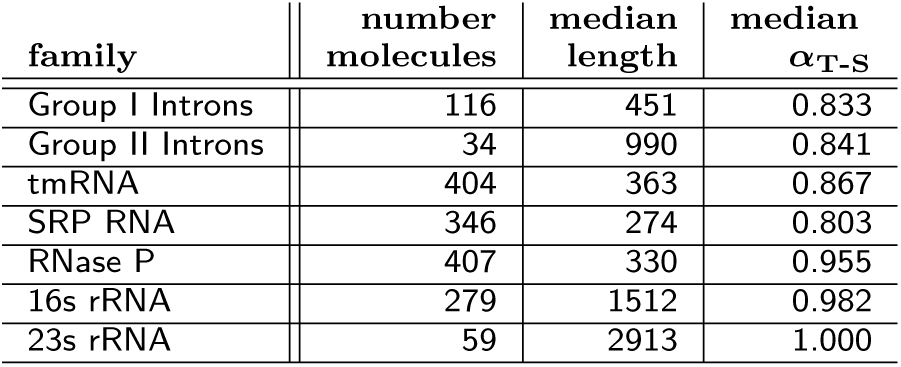
MFE Secondary Structures: calibrated ambiguity indexes, by RNA family. Identical to Table 1, except that the ambiguity indexes and their calibrations are calculated using the MFE secondary structures rather than comparative analyses. There is little evidence in the MFE secondary structures for lower ambiguity indexes among the unbound RNA molecules.

### Formal Statistical Analyses

The T-S ambiguity index *d*_T-S_(*p, s*) is an intra-molecular measure of the difference between the number of available double-stranded Watson-Crick and wobble pairings for segments in and around stems and pseudoknots versus segments within single-stranded regions. As such, *d*_T-S_ depends on both *p* and any purported secondary structure, *s*. Based on a calibrated version, *α*_T-S_(*p, s*), and employing the comparative secondary structure for *s*, we found support for the idea that non-coding RNA molecules in the unbound families, which are active absent participation in ribonucleoproteins, are more likely to have small ambiguity indexes than RNA molecules that operate exclusively as part of ribonucleoproteins. Furthermore, the difference appears to be sensitive to the approach used for identifying secondary structure—there is little, if any, evidence in indexes *d*_T-S_ derived from the MFE secondary structures for lower ambiguities among unbound molecules.

These qualitative observations can be used to formulate precise statistical hypothesis tests. Many tests come to mind, but perhaps the simplest and most transparent are based on nothing more than the molecule-by-molecule signs of the ambiguity indexes. Whereas ignoring the actual values of the indexes is inefficient in terms of information, and probably also in the strict statistical sense, tests based on signs require very few assumptions and are, therefore, more robust to model misspecification. All of the p-values that we will report are based on the hypergeometric distribution, which arises as follows.

We are given a population of *M* molecules, *m* = 1, *…, M*, each with a binary outcome measure *B*_*m*_ *∈ {−*1, +1*}*. There are two subpopulations of interest: the first *M*_1_ molecules make up population 1 and the next *M*_2_ molecules make up population 2; *M*_1_ + *M*_2_ = *M*. We observe *n*_1_ plus values in population 1 and *n*_2_ in population 2

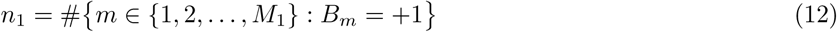

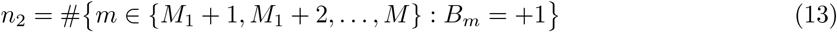

We suspect that population 1 has less than its share of plus ones, meaning that the *n*_1_ + *n*_2_ population of plus ones was not randomly distributed among the *M* molecules. To be precise, let *N* be the number of plus ones that appear from a draw, without replacement, of *M*_1_ samples from *B*_1_, *…, B*_*M*_. Under the null hypothesis, *H*_*o*_, *n*_1_ is a sample from the hypergeometric distribution on *N*:

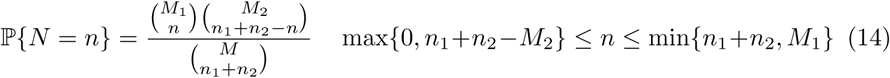

The alternative hypothesis, *H*_*a*_, is that *n*_1_ is too small to be consistent with *H*_*o*_, leading to a left-tail test with p-value ℙ*{N ≤ n*_1_*}* (which can be computed directly or using a statistical package, e.g. *hypergeom.cdf* in *scipy.stats*).

It is by now well recognized that p-values should never be the end of the story. One reason is that *any* departure from the null hypothesis in the direction of the alternative, no matter how small, is doomed to be statistically significant, with arbitrarily small p-value, once the sample size is sufficiently large. In other words, the effect size remains hidden. Therefore, in addition to reporting p-values, we will also display estimated ROC curves, summarizing performance of two related classification problems: (i) Classify a single RNA molecule, randomly selected from the seven families, as belonging to the unbound group or the bound group based *only* on thresholding *d*_T-S_(*p, s*). Compare performance under each of the two secondary-structure models, comparative and MFE; and (ii) Randomly select an RNA molecule from the unbound group and classify the origin of its secondary structure (comparative or MFE), here again based only on thresholding *d*_T-S_(*p, s*). Now Repeat the process, but selecting randomly from the bound group.

### Bound versus Unbound

#### Classification

Consider an RNA molecule, *m*, selected from one of the seven families in our data set, with primary structure *p* and secondary structure *s* computed by comparative analysis. Given only the T-S ambiguity index of *m* (i.e. given only *d*_T-S_(*p, s*)), how accurately could we classify the origin of *m* as the unbound versus bound group? The foregoing exploratory analysis suggests constructing a classifier that declares a molecule to be unbound when *d*_T-S_(*p, s*) is small, e.g. *d*_T-S_(*p, s*) *< t*, where the threshold *t* governs the familiar trade off between rates of “true positives” (an unbound molecule *m* is declared ‘unbound’) and “false positives” (a bound molecule *m* is declared ‘unbound’). Small values of *t* favor low rates of false positives at the price of low rates of true positives, whereas large values of *t* favor high rates of true-positives at the price of high rates of false positives. Since for each molecule *m* we have both the correct classification (unbound or bound) and the statistic *d*, we can estimate the ROC performance of our threshold classifier by plotting the empirical values of the pair

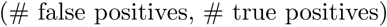

for each value of *t*. The ROC curve for the two-category (unbound versus bound) classifier based on thresholding *d*_T-S_(*p, s*) *< t* is shown in the left panel of Figure 1. Also shown is the estimated area under the curve (AUC=0.81), which has a convenient and intuitive interpretation, as it is equal to the probability that for two randomly selected molecules, *m* from the unbound population and *m*′ from the bound population, the T-S ambiguity index of *m* will be smaller than the T-S ambiguity index of *m*′.

**Figure 1.**
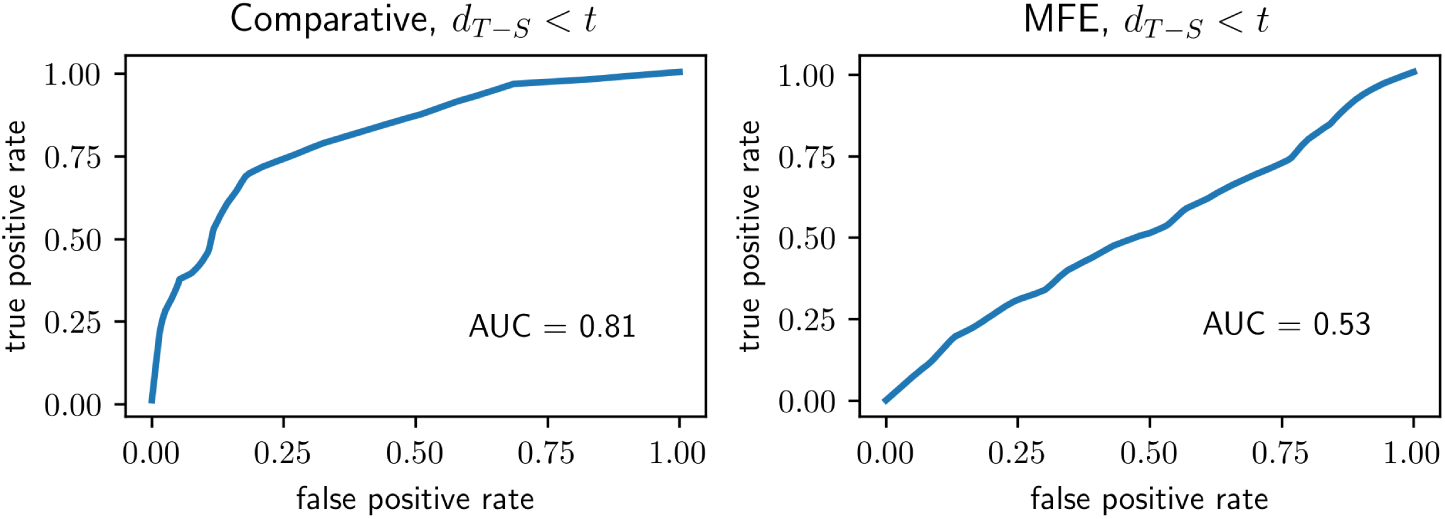
Unbound or Bound? ROC performance of classifiers based on thresholding the T-S ambiguity index. Small values of *d*_T-S_(*p, s*) are taken as evidence that a molecule belongs to the unbound group as opposed to the bound group. In the left panel, the classifier is based on using the comparative secondary structure for *s* to compute the ambiguity index. Alternatively, the MFE structure is used for the classifier depicted in the right panel. *AUC:* Area Under Curve—see text for interpretation. Additionally, for each of the two experiments, a p-value was calculated based only on the signs of the individual ambiguity indexes, under the null hypothesis that positive indexes are distributed randomly among molecules in all seven RNA families. Under the alternative, positive indexes are more typically found among the unbound as opposed to bound families. Under the null hypothesis the test statistic is hypergeometric—see Eq 14. *Left Panel*: *p* = 1.2 *×* 10^*−*34^. *Right Panel*: *p* = 0.02. In considering these p-values, it is worth re-emphasizing the points made about the interpretation of p-values in the paragraph following Eq 14. The right panel illustrates the point: the ambiguity index based on the MFE secondary structure “significantly distinguishes the two categories (*p* = 0.02)” but clearly has no utility for classification. (These ROC curves and those in Figure 2 were lightly smoothed by the method known as “Locally Weighted Scatterplot Smoothing,” e.g. with the python command Y=lowess(Y, X, 0.1, return sorted=False) coming from *statsmodels.nonparametric.smoothers lowess*.)

#### p-Values

As mentioned earlier, we can also associate a traditional p-value to the problem of separating unbound from bound molecules, based again on the T-S ambiguity indexes. We consider only the signs (positive or negative) of these indexes, and then test whether there are fewer than expected positive indexes among the unbound as opposed to the bound populations. This amounts to computing ℙ*{N ≤ n*_1_*}* from the hypergeometric distribution—Eq (14). The relevant statistics can be found in Table 3, under the column labels **#mol’s** and #***d***_**T-S**_ ***>* 0**. Specifically, *M*_1_ = 116 + 34 = 150 (number of unbound molecules), *M*_2_ = 404 + 346 + 407 + 279 + 59 = 1495 (number of bound molecules), *n*_1_ = 50 + 8 = 58 (number of positive T-S indexes among unbound molecules) and *n*_2_ = 368 + 269 + 379 + 210 + 53 = 1279 (positive bound indexes). The resulting p-value, 1.2·10^*−*34^, is essentially zero, meaning that the positive T-S indexes are not distributed proportional to the sizes of the unbound and bound populations, which is by now obvious in any case. To repeat our caution, small p-values conflate sample size with effect size, and for that reason we have chosen additional ways, using permutations as well as classifications, to look at the data.

**Table 3.** Numbers of Positive Ambiguity Indexes, by family. #mol’s: number of molecules; **#*d***_**T-S**_ ***>* 0**: numbers of positive T-S ambiguity indexes, secondary structures computed by *comparative analysis*; 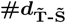numbers of positive T-S ambiguity indexes, secondary structures computed by *minimum free energy*.

| family | #mol's | $\#d_{T-S} > 0$ | $\#d_{\tilde{T}-\tilde{S}} > 0$ |
| --- | --- | --- | --- |
| Group I Introns | 116 | 50 | 94 |
| Group II Introns | 34 | 8 | 27 |
| tmRNA | 404 | 368 | 358 |
| SRP RNA | 346 | 269 | 264 |
| RNase P | 407 | 379 | 377 |
| 16s rRNA | 279 | 210 | 254 |
| 23s rRNA | 59 | 53 | 54 |

### Comparative versus Minimum Free Energy

As we have just seen, ambiguity indexes based on MFE secondary structures, as opposed to comparative secondary structures, do not make the same stark distinction between unbound and bound RNA molecules. To explore this a little further, we can turn the analyses of the previous paragraphs around and ask to what extent knowledge of the ambiguity index is sufficient to predict the source of a secondary structure—comparative or free energy? This turns out to depend on the group from which the molecule was drawn: The ambiguity index is strongly predictive among unbound molecules and, at best, weakly predictive among bound molecules.

Consider the two ROC curves in Figure 2. In each of the two experiments a classifier was constructed by thresholding the T-S ambiguity index, declaring the secondary structure, *s*, to be “comparative” when *d*_T-S_(*p, s*) *< t* and “MFE” otherwise. The difference between the two panels is in the population used for the classification experiments—unbound molecules in the left-hand panel (AUC=0.81) and bound molecules in the right-hand panel (AUC=0.54, barely above chance). The corresponding hypothesis tests seek evidence against the null hypotheses that in a given group (unbound or bound) the set of positive T-S ambiguity indexes (*d*_T-S_(*p, s*) *>* 0) are equally distributed between the comparative and free-energy derived indexes, and in favor of the alternatives that the T-S ambiguity indexes are less typically positive for the comparative secondary structures. The necessary data can be found in Table 3. The test results are consistent with the classification experiments: the hypergeometric p-value is is 5.4 · 10^*−*14^ for the unbound population and 0.07 for the bound population.

**Figure 2.**
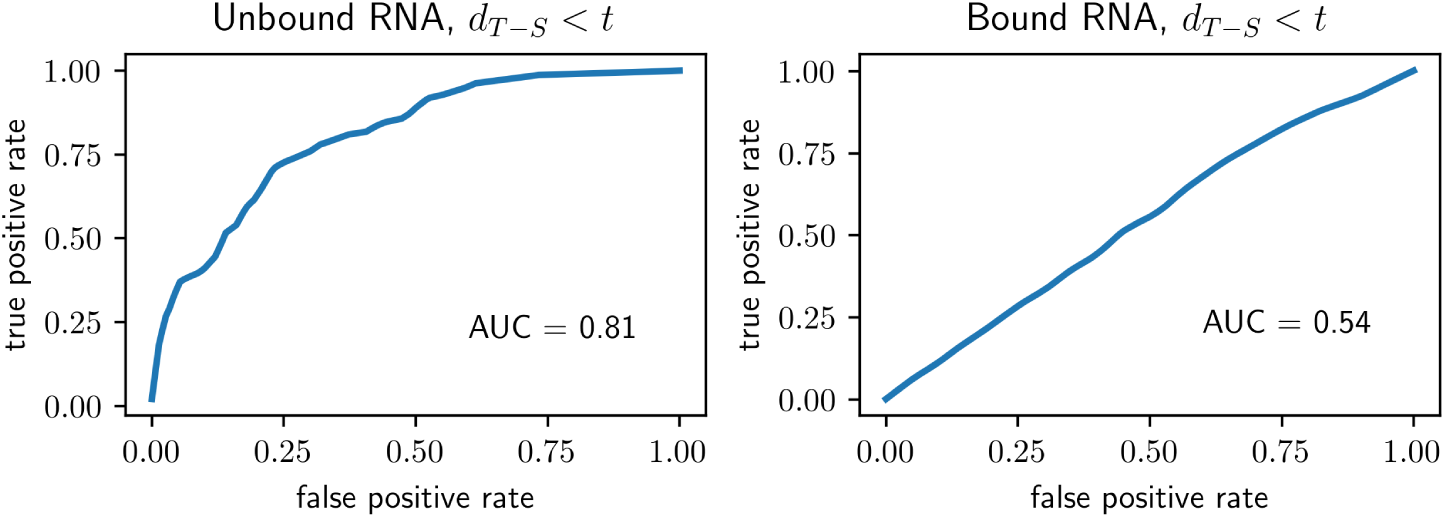
Comparative or MFE? As in Figure 1, each panel depicts the ROC performance of a classifier based on thresholding the T-S ambiguity index, with small values of *d*_T-S_(*p, s*) taken as evidence that *s* was derived by comparative as opposed to MFE secondary structure analysis. *Left Panel*: performance on molecules chosen from the unbound group. *Right Panel*: performance on molecules chosen from the bound group. Conditional p-values were also calculated, using the hypergeometric distribution and based only on the signs of the indexes. In each case the null hypothesis is that comparative secondary structures are as likely to lead to positive ambiguity indexes as are MFE structures, whereas the alternative is that positive ambiguity indexes are more typical when derived from MFE structures. *Left Panel*: *p* = 5.4 *×* 10^*−*14^. *Right Panel*: *p* = 0.07.

Qualitatively, these various ROC and p-value results were easy to anticipate from even a superficial examination of Table 3. Start with the first two rows (unbound molecules): A relatively small fraction of unbound molecules have positive ambiguities when the index is computed from comparative analyses, whereas most of these same molecules have positive ambiguities when the index is computed from MFE structures. Looking across the next five rows (bound molecules), no such trend is discernible. Similarly, from a glance at the column labeled #*d*_T-S_ *>* 0 (derived from comparative analyses) it is apparent that the fraction of positive indexes among the unbound molecules is much lower than among the bound molecules. What’s more, this effect is missing in the MFE indexes 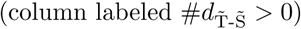.^[4]^

## Discussions and Conclusions

Consider a non-coding RNA molecule with a native tertiary structure that is active, *in vivo*, without necessarily being tightly bound with other molecules in a ribonucleoprotein complex. We have labeled these molecules “unbound” and reasoned that there are likely relationships between their primary and secondary structures that not only support the tertiary structure, but also the folding process by which it emerges. Specifically, we reasoned that examination of the primary and native secondary structures might reveal evolutionary mechanisms that discourage disruptive kinetic traps. Conjecturing that the availability of non-native pairings for subsequences that are part of the native secondary structure would be particularly disruptive, we defined an intra-molecular index that we called the ambiguity index. The ambiguity index is a function of a molecule’s primary and native secondary structures devised so that lower values of the index reflect fewer opportunities for stem participating subsequences to pair elsewhere in the molecule. We examined the Group I and Group II introns, two families of molecules that are believed to perform some of their functions (namely self splicing) in an “unbound” state, to see if their ambiguity indexes were lower than might be expected were there no such evolutionary pressures to protect stem structures. Heuristic permutation-type tests appeared to confirm our expectation that these molecules would have low ambiguities.

We sought additional evidence in two directions. The first was to compare ambiguity indexes in unbound molecules to those in “bound” molecules, i.e. molecules that are known to function as part of ribonucleoprotein complexes where the argument against these particular kinds of ambiguities is weaker. We found a strong separation between the unbound and bound molecules, the former having substantially lower indexes. This was demonstrated by statistical tests and, perhaps more meaningfully, by showing that the ambiguity index could be used to classify with good accuracy individual molecules as either bound or unbound. These experiments were based on comparative secondary structure analysis, which remains the most trusted source for most RNA molecules [12, 13].

In a second approach to additional evidence we substituted the comparative secondary structures with ones that were derived from approximations to the thermodynamic equilibrium structure (minimum free energy—“MFE” structures). Though less accurate, MFE and related equilibrium-type structures are easy and quick to compute. But one line of thinking is that active biological structures are determined more by kinetic accessibility than thermodynamic equilibrium per se. Biological stability is relative to biological timescale; the folding of any particular RNA could just as well end in metastability, provided that the process is repeatable and the result sufficiently stable over the molecule’s proper biological lifetime. Indeed, it would be arguably easier to evolve an effective tertiary structure without the additional and unnecessary burden of thermal equilibrium. To the extent that kinetic accessibility and metastability might be more relevant than thermodynamic equilibrium, there would be little reason to expect the ambiguity index to make the same separation between unbound and bound molecules when derived from MFE structures instead of comparative structures. The results were consistent with this point of view—ambiguity indexes based on MFE structures make weak classifiers. We were surprised by the strength of the effect. After all, MFE structures are superficially quite similar to comparative structures, yet the classification performance goes from strong (*>* 80% AUC) to negligible (53% AUC, just above chance). A worthwhile follow-up would be to examine the actual differences in secondary structure (as was done, with similar motivation but different tools, in [20]) in an effort to discern how they impact ambiguity.

However, and this has often been pointed out (e.g. [28, 29]), the MFE structure itself may be a poor representative of thermal equilibrium. It is possible, then, that our observations to the effect that comparative and MFE structures have substantially different relationships to the ambiguity indexes, and our interpretation that comparative structures better separate unbound from bound molecules, would not hold up as well if we were to adopt a more ensemble-oriented structure in place of the MFE, as advocated by [30], for example. In a related vein, and also within the context of thermodynamic equilibrium, Lin et al. [31] have given evidence that competing stems which are inconsistent may *both* contain a high measure of information about the equilibrium distribution, suggesting that in such cases both forms could be active and the notion of *single* (locations we have labeled “S”) might itself be ambiguous. Certainly there are RNA molecules that are active in more than one structural conformation.

The ambiguity index *d*_T-S_ is derived from the difference in average ambiguities of subsequences partly paired in the native structure (“T”, transition locations) from those not paired in the native structure (single locations). We expected these differences to be small in unbound as opposed to bound molecules because we expected the stem structures to be more protected from non-native pairings. But this coin has another side: low ambiguities at unpaired (single) locations of *bound* molecules relative to unbound molecules would have the same effect. As an example, some unpaired RNA sequences may be critical to function, as in the messenger RNA-like (“MLR”) region of tmRNA, and therefore relatively unambiguous. Also, it is possible that the formation of non-native stems among single-type subsequences are particularly disruptive to, perhaps even stereochemically preventing, the binding of an RNA molecule into a ribonucleoprotein complex. More generally, it is reasonable to assume that different evolutionary forces are at play for molecules destined to operate as parts of ribonucleoprotein complexes. In any case, the folding story may be even more complicated, or at least quite different, for the ribonculeoprotein RNAs.

Finally, we note that the ambiguity index, as currently formulated, is symmetric in the sense that there is no explicit difference in contributions from different locations along the 5′ to 3′ axis. Yet cotranscriptional folding, which appears to be nearly universal in non-coding RNA [32] strongly suggests that not all ambiguities are equally disruptive. Indeed, some non-native pairings between two subsequences, one of which is near the 3′ end of the molecule, might have been rendered stereochemically impossible before the 3′ half has even been transcribed. Cotranscriptional folding opens up many such lines of reasoning, most of which suggest new indexes that could be statistically explored, especially as the data bank of known structures and functions continues to grow.

Overall, our results are consistent in supporting a role for kinetic accessibility that is already visible in the relationship between primary and secondary structures. Stronger evidence will require more bound and unbound families. The limiting factors, as of today, are the availability of families with large RNA molecules for which the comparative structures have been worked out and largely agreed upon.

## Methods

### Datasets

We obtained comparative-analysis secondary structure data for seven different families of RNA molecules from the RNA STRAND database [33], a curated collection of RNA secondary structures. These families include: Group I Introns and Group II Introns [34], tmRNAs and SRP RNAs [35], the Ribonuclease P RNAs [36], and 16s rRNAs and 23s rRNAs [34]. Table 4 contains information about the numbers and lengths (measured in nucleotides) of the RNA molecules in each of the seven families. Note that we excluded families like tRNAs, 5s rRNAs and hammerhead ribozymes since most of the molecules in these families are too short to be of interest for our purpose. Also, since we are focusing on comparative-analysis secondary structures, to be consistent, we excluded any secondary structures derived from X-ray crystallography or NMR structures.

**Table 4.** Data Summary. The seven families of RNA used in the experiments. Table includes the number of molecules in each family, as well as basic statistics about the numbers of nucleotides in the primary sequence of each of the molecules. Data was downloaded from the RNA STRAND database.

| family | number | min length | max length | median |
| --- | --- | --- | --- | --- |
| Group I Introns | 116 | 210 | 2630 | 451 |
| Group II Introns | 34 | 619 | 2729 | 990 |
| tmRNA | 404 | 102 | 437 | 363 |
| SRP RNA | 346 | 66 | 533 | 274 |
| RNase P | 407 | 189 | 486 | 330 |
| 16s rRNA | 279 | 612 | 2394 | 1512 |
| 23s rRNA | 59 | 953 | 4381 | 2913 |

Note that Group I and Group II Introns are the only available families of unbound RNAs suitable for our analysis. There are some other families of unbound RNAs (e.g. ribozymes), but most of these RNAs are too short in length, and many of the structures are not derived using comparative analysis. Hence they are not included.

### RNA Secondary Structure Prediction Methods

Comparative analysis [37] is based on the simple principle that a single RNA secondary structure can be formed from different RNA sequences. Using alignments of homologous sequences, comparative analysis has proven to be highly accurate in determining RNA secondary structures [13]. We used a large set of RNA secondary structures determined by comparative analyses to serve as ground truth.

When it comes to computational prediction of RNA secondary structures, exact dynamic programming algorithms based on carefully measured thermodynamic parameters make up the most prevalent methods. There exist a large number of software packages for the energy minimization [28, 38–43]. In this paper, we used the ViennaRNA package [38] to obtain the MFE secondary structures for our statistical analysis.

### Reproducing the Results

The results presented in this paper can be easily reproduced. Follow the instructions on https://github.com/StannisZhou/rna_statistics. Here we make a few comments regarding some implementation details.

- In the process of obtaining the data, we used the *bpseq* format, and excluded structures derived from X-ray crystallography or NMR structures, as well as structures for duplicate sequences. Concretely, this means picking a particular type, and select *No* for *Validated by NMR or X-Ray* and *Non-redundant sequences only* for *Duplicates* on the search page of the RNA STRAND database. A copy of the data we used is included in the GitHub repository, but the same analyses can be easily applied to other data.
- When processing the data, we ignored molecules for which we have nucleotides other than *A, G, C, U*, and molecules for which we don’t have any base pairs.
- When comparing the local ambiguities in different regions of the RNA molecules, we ignored molecules for which we have empty regions (i.e. at least one of *single, double* and *transitional* is empty), as well as molecules where all local ambiguities in *single* or *double* regions are 0.
- For shuffling primary structures, we used an efficient and flexible implementation of the Euler algorithm [25–27] called uShuffle [44], which is conveniently available as a python package.

## Competing interests

The authors declare that they have no competing interests.

**Author’s contributions**

## Acknowledgements

The authors would like to thank Yang Chen for helpful discussions and Matthew T. Harrison and Charles Lawrence for many valuable suggestions.

## Footnotes

[1] By which we will mean sequences of G·U (“wobble pairs”) and/or Watson-Crick pairs.

[2] Native secondary structures often include so-called pseudoknots, which are some-times excluded, or handled separately, for computational efficiency. Pseudoknots are formed from paired complementary subsequences and therefore included, by definition, in the ambiguity index.

[3] Molecular dynamics, which might be called “agnostic” to the question of equilibrium, has proven to be exceedingly difficult, and has not yet yielded a useful tool for generic folding of large molecules.

[4] The specific values of the areas under the ROC curves depend on the specific values of the indexes. The equality—to two digits—of the areas in the left-hand panels of Figures 2 and 1 is a coincidence.

